# Expression from DIF1-motif promoters of *hetR* and *patS* is dependent on HetZ and modulated by PatU3 during heterocyst differentiation

**DOI:** 10.1101/2020.04.15.042648

**Authors:** Yaru Du, He Zhang, Hong Wang, Shuai Wang, Qiqin Lei, Chao Li, Renqiu Kong, Xudong Xu

## Abstract

HetR and PatS/PatX-derived peptides are the activator and diffusible inhibitor for cell differentiation and patterning in heterocyst-forming cyanobacteria. HetR regulates target genes via HetR-recognition sites. However, some genes (such as *patS/patX*) upregulated at the early stage of heterocyst differentiation possess DIF1 (or DIF^+^) motif (TCCGGA) promoters rather than HetR-recognition sites; *hetR* possesses both regulatory elements. How HetR controls heterocyst-specific expression from DIF1 motif promoters remains to be answered. This study presents evidence that the expression from DIF1 motif promoters of *hetR, patS* and *patX* is more directly dependent on *hetZ*, a gene regulated by HetR. The HetR-binding site upstream of *hetR* is not required for its autoregulation. PatU3 (3’ portion of PatU) that interacts with HetZ may modulate HetZ-dependent gene expression. These findings contribute to understanding of the mutual regulation of *hetR, hetZ-patU* and *patS/patX* in a large group of multicellular cyanobacteria.

## Introduction

Cyanobacteria were the first group of microorganisms that performed oxygenic photosynthesis [1, 2]. In the early earth environment, nitrogen nutrient was a limiting factor for propagation of microbes. Under this selective pressure, *nif* genes spread among bacteria, and some cyanobacteria acquired the N_2_ fixation capability. With the rise of atmospheric oxygen, certain filamentous species developed the capability to form specialized N_2_-fixing cells, called heterocysts, to protect nitrogenase from inactivation by oxygen [3–5]. Nowadays, heterocyst-forming cyanobacteria contribute significantly to nitrogen fixation in the earth’s biosphere [6–8]. In species from different genera of heterocyst-forming cyanobacteria, heterocysts are differentiated at one end, two ends, or intercalary positions of filaments [9]. *Anabaena* sp. PCC 7120 (hereafter *Anabaena* 7120) was derived from a species that produces semi-regularly spaced single heterocysts along non-branched filaments in response to nitrogen stepdown. It is the most often used model strain for molecular studies on heterocyst-related topics [10].

Heterocyst differentiation and pattern formation largely depend on the key regulator HetR [11] and RGSGR-containing peptides, which are derived from PatS [12, 13], PatX [14] or HetN [15], representing an example of the most ancient activator-inhibitor (reaction-diffusion) patterning process [16–18]. In *Anabaena* 7120, PatS is the main source of morphogen for de novo pattern formation [13], while HetN is required for maintenance of the pattern [19]. HetR is the only known target of RGSGR-containing peptides [20], and it binds to consensus recognition sites upstream of *hetP* [21, 22], *hetZ* [23] and several other genes, including its own encoding gene [24–26]. Among these genes, *hetZ* is involved in control of heterocyst differentiation at an early stage [27], and *hetP* is required for commitment to heterocyst differentiation [28]. *hetZ* and *hetP* functionally overlap with each other, and co-expression of these two genes was shown to restore heterocyst formation in *hetR*-null mutants [29]. In a different substrain of *Anabaena* 7120, expression of *hetZ* alone restored heterocyst formation in a *hetR*-deletion mutant [30]. The variable requirement for *hetP* expression may depend on differences in genetic backgrounds of substrains [31]. *hetP* and *hetZ* are both upregulated in differentiating cells, as a result of the accumulation of HetR [21, 23]. *patS* is also upregulated in differentiating cells [12], but no consensus recognition site for HetR is present in the sequence upstream of *patS*.

Immediately downstream of *hetZ* in many filamentous cyanobacteria is a gene called *patU*; these two genes, together with *hetR*, are listed among the core set of genes for filamentous species [27, 32]. In *Anabaena* 7120, *patU* is split into *patU5* and *patU3* [27]. *hetZ* and *patU3* play opposite roles in heterocyst differentiation: *hetZ* promotes, while *patU3* inhibits [27].

Before the consensus HetR-recognition sequence was identified, DIF^+^ (later called DIF1) motif (TCCGGA) had been bioinformatically identified in sequences upstream of *hetR* and several other genes in *Anabaena* 7120 [33]. More recently, the DIF1 motif was proposed as a consensus regulatory sequence (centered at −35 region) for *patS* and *patX* in heterocyst-forming cyanobacteria [34]. The role of DIF1 motif in expression of the *nsiR* promoter [33] and a synthetic minimal promoter has been reported [35]. However, the role of predicted DIF1 motif promoters in expression of *hetR*, *patS* and *patX* has not been shown experimentally. In particular, HetR-recognition site and DIF1 motif are both present upstream of *hetR*. Most importantly, which of HetR, HetZ and HetP is required for the regulation of DIF1-motif promoters? In *Anabaena* 7120, deletion of *hetZ* blocked the induced expression of *hetR*, *hetP* and *patS*, whereas *hetP* showed no effects on these genes [30]. This result excluded HetP, but did not establish either HetR or HetZ, as the factor required for the induced expression of *hetR* and *patS*. More generally, the expression from DIF1 motif promoters is dependent on a functional *hetR* [33, 36]. In this study, we found that HetZ is more directly involved in the regulation of DIF1-motif promoters of *hetR* and *patS*. In addition, PatU3 that interacts with HetZ may modulate the expression of these genes.

## Materials and methods

### General

*Anabaena* 7120 and derivatives, listed in Table S1, were cultured in BG11 medium in the light of 30 μE m^−2^ s^−1^ on a rotary shaker. Erythromycin (5 μg ml^−1^), neomycin (20 μg ml^−1^) or spectinomycin (10 μg ml^−1^) was added to the medium as appropriate. For nitrogen stepdown, *Anabaena* 7120 grown in BG11 (OD_730_, 0.7~0.9) was collected by centrifugation, washed 3 times with BG110 (without nitrate) and resuspended in the same medium for indicated hours. Microscopy was performed as previously described [38].

### Construction of plasmids and *Anabaena* strains

Plasmid construction processes are described in Table S1 in the supplemental materials. DNA fragments cloned by PCR were confirmed by sequencing.

Plasmids were introduced into *Anabaena* 7120 and mutants by conjugation [48]. Homologous double-crossover recombinants were generated based on positive selection with *sacB* [49]. The complete segregation of mutants was confirmed by PCR. *Anabaena* strains and primers are listed in Table S1.

### Transcription analyses

RNA extraction, elimination of residual DNA and reverse transcriptase quantitative polymerase chain reaction (RT-qPCR) were performed as we described before [29]. PCR primers (indicated with ‘RT’ in name) are listed in Table S1.

Promoter activities were visualized using *gfp* (green fluorescence protein) as the reporter gene. Relative copy numbers of zeta- or pDU1-based plasmids (relative to *rnpB*) were evaluated by quantitative PCR as described in the reference [26] using primers gfp-1/gfp-2, pDU1-1/pDU1-2 and rnpB-1/rnpB-2 listed in Table S1.

### Rapid amplification of cDNA ends (RACE)

RACE was performed according to Zhang et al. [27], using universal primer/hetR-race-1 and nested universal primer/hetR-race-2 as the primers for 2 rounds of PCR. The universal primer and nested universal primer were provided with the SMART RACE cDNA amplification kit (Clontech, TaKaRa Bio., Otsu, Japan); hetR-race-1 and hetR-race-2 are listed in Table S1. Transcription start points were determined based on sequencing of RACE products. Two biological repeats showed similar results.

### Western blot analysis

*Anabaena* 7120 was deprived of fixed nitrogen for 24 h, harvested by centrifugation, washed with 20 mM Tris-HCl (pH 8.0) containing 1 mM PMSF and resuspended in the same buffer. Cells were broken with a French press (SCIENTZ, China) at 240 MPa (cell pressure) and centrifuged at 12,000 × g for 15 min. The supernatant was used as cell extracts for the Western blot analysis.

Proteins were separated by 12% SDS-PAGE and electro-blotted onto NC filters. HetR and HetZ were detected with rabbit antiserum against purified HetR or HetZ overproduced in *Escherichia coli*, visualized using alkaline phosphatase-conjugated secondary antibody specific for rabbit IgG (Thermo Scientific, USA) with NBT and BCIP as substrates. Two biological repeats showed similar results.

## Results

### Upregulated expression from P*_hetR_* and P_*patS*_ in *hetR*-minus heterocysts

In a *hetR*-minus mutant, heterocyst differentiation is not initiated, and genes otherwise specifically expressed in heterocysts are mostly not upregulated after nitrogen stepdown. Such genes could be directly or indirectly regulated by HetR. Under our conditions, co-expression of *hetZ* and *hetP* from P*_ntcA_* (rather than expression of *hetZ* or *hetP* alone from the same promoter) enabled the *hetR* mutant, 7120*hetR*::C.CE2, to form functional heterocysts at the ends of filaments [29]. Such a phenotype was probably due to the lack of expression of *patA*, a gene required for heterocyst formation at intercalary positions, in vegetative cells of the *hetR* mutant [26]. Formation of functional *hetR*-minus heterocysts [29] indicated that genes required for the function of heterocysts are properly expressed but not necessarily that P*_hetR_* and P*_patS_* are upregulated in (pro)heterocysts. This system allowed us to test the promoters of *hetR* and *patS* in (pro)heterocysts without the presence of HetR.

Plasmids carrying P*_ntcA_-hetZ-hetP* and the structure ‘Ω-promoter-*gfp*’ (the Ω cassette terminates background transcription, ref. 37; *gfp*, green fluorescence protein) were constructed and introduced into the *hetR* mutant. The tested promoters included P_*hetR*_, P*_patS_*, P*_hepB_* and P*_hglD_*. *hepB* and *hglD* are involved in the formation of heterocyst envelope polysaccharide layer and glycolipid layer respectively, therefore P*_hepB_* and P*_hglD_* were included as the controls for heterocyst-specific expression [38]. Without a functional *hetR*, overexpression of *hetZ* and *hetP* led to heterocyst formation at the ends of filaments and upregulated expression of *gfp* from P_*hetR*_, P_*patS*_, P*_hepB_* and P*_hglD_* in heterocysts relative to vegetative cells (Fig. 1). Clearly, HetR is not essential for the expression from all these promoters.

**Fig 1.**
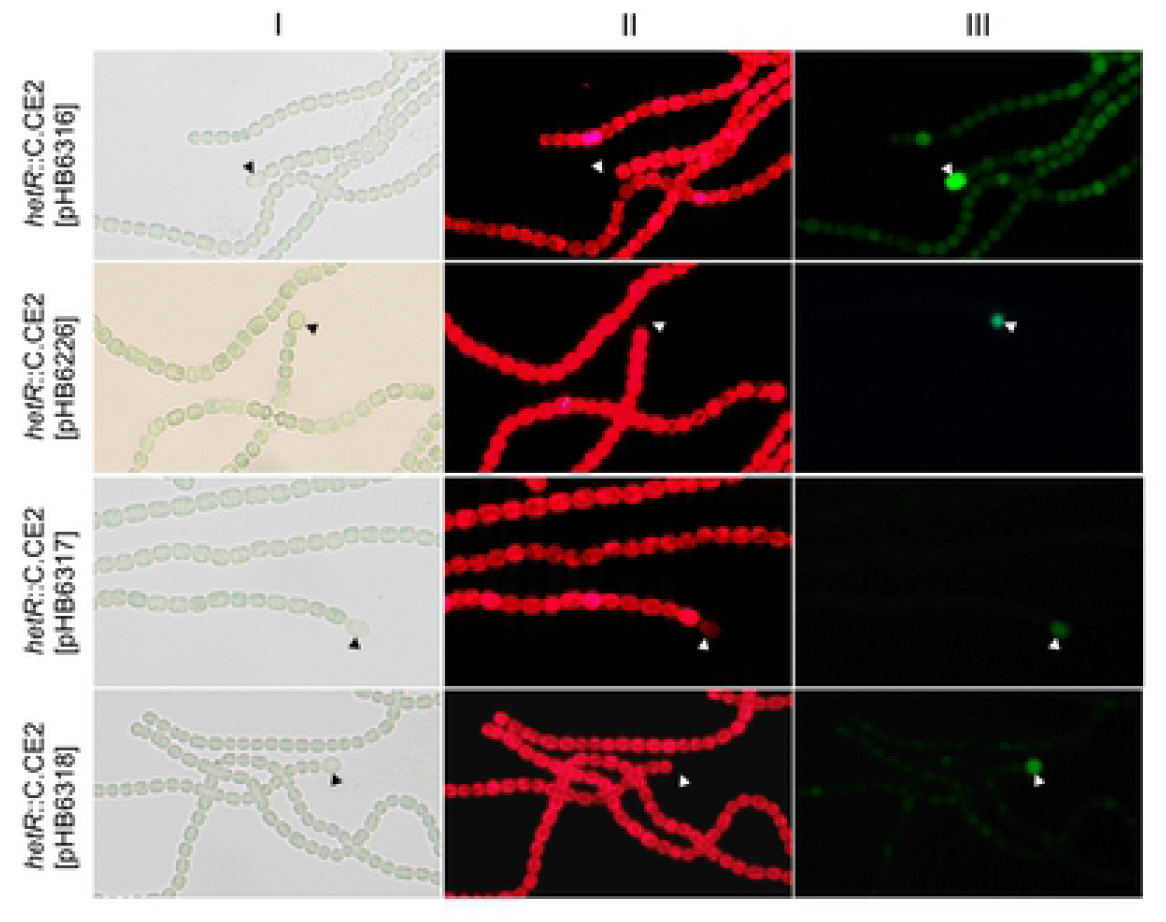
Light (I), autofluorescence (II) and GFP fluorescence (III) photomicrographs showing the expression of *gfp* from the promoter of *hetR*, *patS*, *hepB* or *hglD* in *hetR*-minus heterocysts. Plasmids with P*_ntcA_-hetP-hetZ* and Ω-promoter-*gfp* (P*_hetR_*, P*_patS_*, P*_hepB_* and P*_hglD_* carried on pHB6316, pHB6226, pHB6317 and pHB6318 respectively) were introduced into *Anabaena* 7120 *hetR*::C.CE2. Solid arrowheads point to heterocysts.

### Upregulation of *patS* in heterocysts depends on the DIF1 motif and *hetZ*

In the previous study [29], we examined the expression of several genes in *Anabaena* 7120 and *hetZ*, *hetP* mutants. Using the same mRNA samples, we also performed RT-qPCR analysis of *patS*. The expression of *patS* was shown to be dependent on *hetZ* rather than *hetP* (S1 Fig). Consistently, *patS* was upregulated in a Δ*hetP* mutant but not in a Δ*hetZ* mutant [30].

To confirm the role of HetZ in expression of *patS*, we further generated a partial deletion mutant, 7120*hetZ*del4-201, of *Anabaena* 7120 with 66 amino acids near the N-terminus of HetZ deleted in frame while preserving the putative promoter internal to *hetZ* serving *patU5-patU3* [27]. This mutant showed no morphologically discernible heterocyst differentiation but formed some cells with less autofluorescence after nitrogen stepdown. These cells initiated differentiation, but the differentiation process ceased at the very early stage. The decreased autofluorescence is due to the degradation of phycobilisomes [39]. A non-replicative plasmid (pHB6069) containing P*_patS_* (−1070 ~ +48 relative to the translational start site of *patS*) upstream of *gfp* was integrated into the genomes of *Anabaena* 7120 and the derivative strain 7120*hetZ*del4-201 via homologous single-crossover recombination. *Anabaena* 7120::pHB6069 showed moderate expression of *gfp* specifically in (pro)heterocysts, whereas 7120*hetZ*del4-201::pHB6069 showed much weaker (but visible) expression of *gfp* in differentiating cells (Fig. 2A).

**Fig 2.**
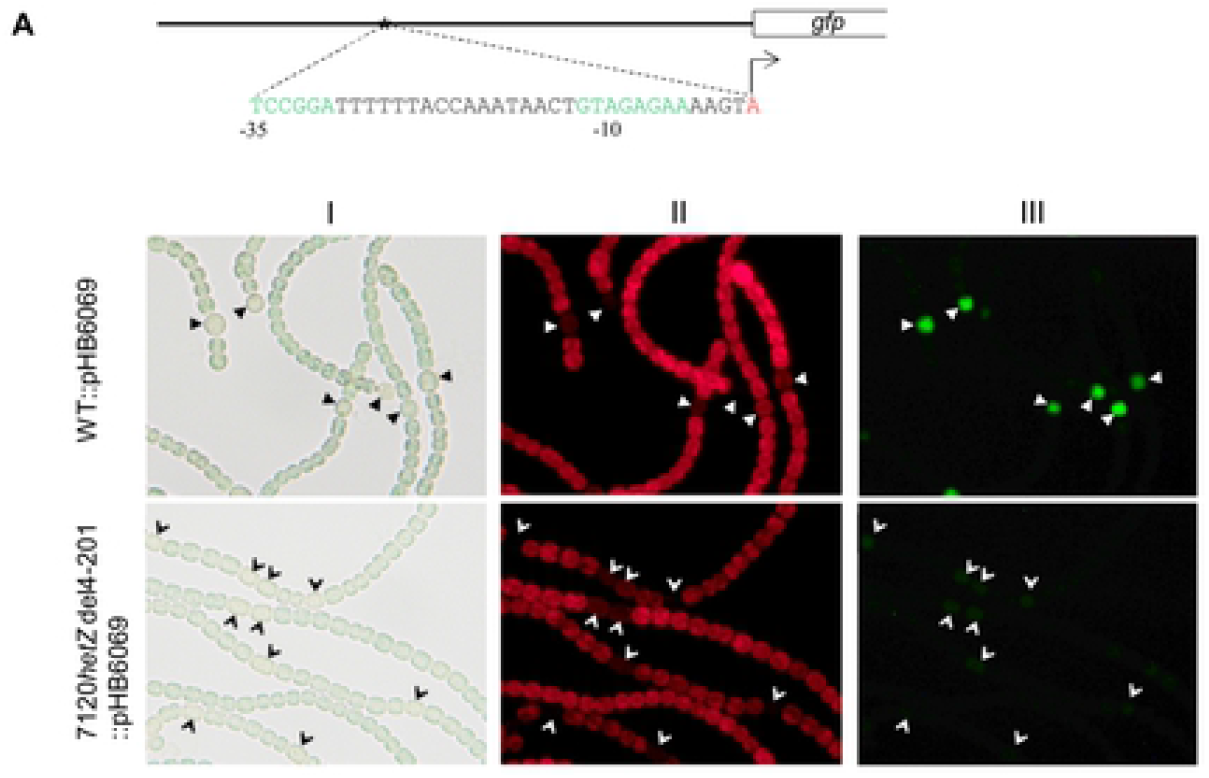
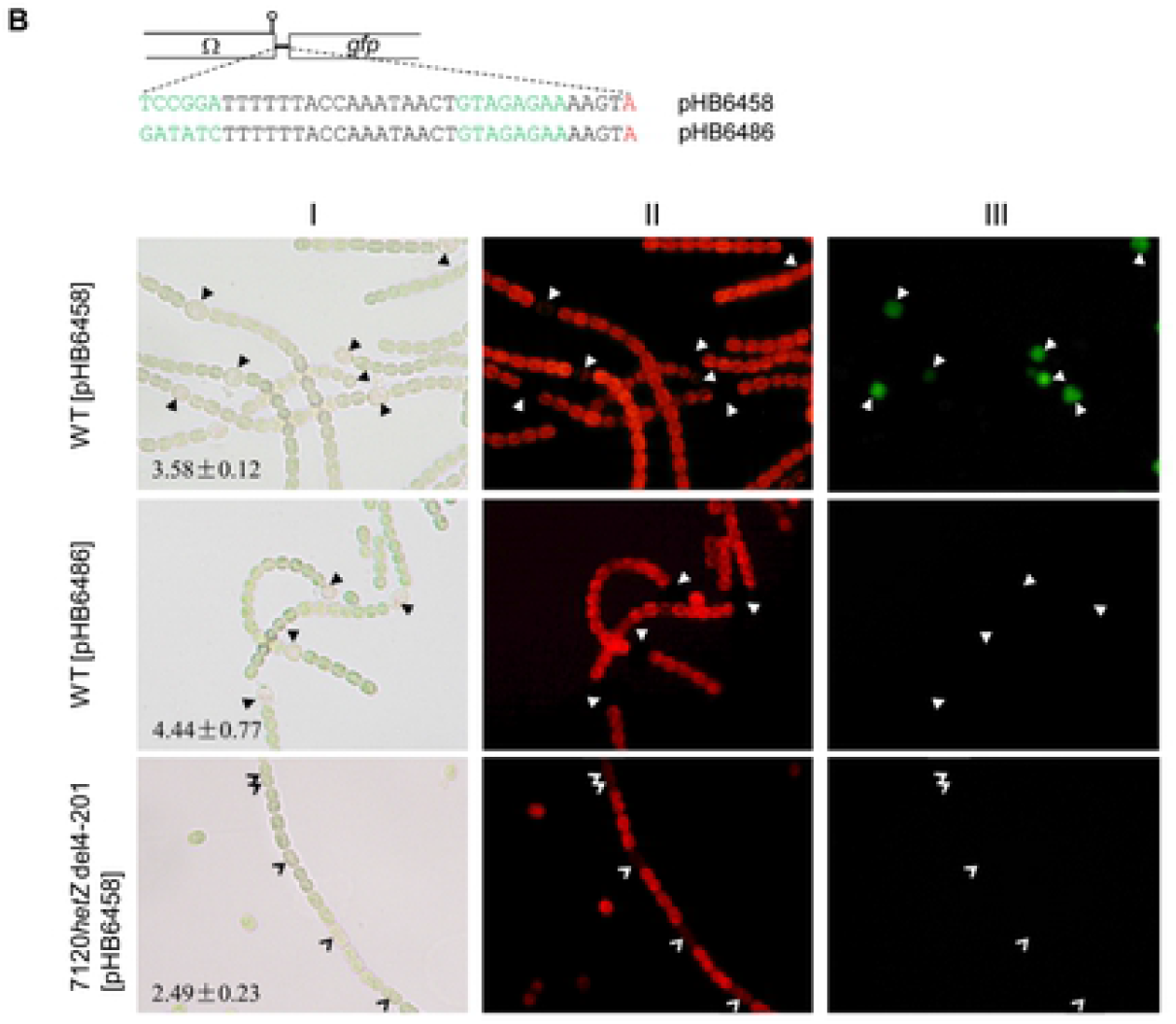
Expression of *gfp* from the *patS* promoter in *Anabaena* 7120 and 7120*hetZ*del4-201. Light (I), autofluorescence (II) and GFP fluorescence (III) photomicrographs of *Anabaena* derivative strains were taken at 24 h after nitrogen stepdown. Solid and empty arrowheads point to heterocysts and differentiating cells. Means ± SD are relative copy numbers of plasmids (relative to the copy number of *rnpB* in the genome). (A) Expression of *gfp* from the full-length *patS* promoter in the genome. The plasmid pHB6069 with P*_patS_-gfp* was integrated into the chromosome of *Anabaena* 7120 and the *hetZ* mutant via homologous single-crossover recombination. In the schematic diagram for the structure of full-length P*_patS_* fused to *gfp*, the bent line with an empty arrowhead indicates the transcription start point of the DIF1-motif promoter. (B) Expression of *gfp* from the minimal DIF1-motif promoter on zeta-based plasmids in *Anabaena* 7120 and the *hetZ* mutant. pHB6486 and pHB6458 are plasmids with the minimal DIF1-motif promoter of *patS*, with TCCGGA substituted or not. The stem-loop structure stands for the transcription terminator at the end of Ω cassette.

Employing *gfp* as a reporter gene in *Anabaena* 7120, we delimited the promoter of *patS* to the region −662 ~ −457 upstream of the start codon (S2 Fig, see photomicrographs for expression of *gfp* from fragments i, ii and iii). In this region, there is a DIF1 motif (TCCGGA) located 35 bp upstream of the tsp (transcriptional start point) −580 of *patS* [34]. We constructed a zeta-based plasmid with the minimal DIF1-motif promoter (a 41-bp fragment) positioned upstream of *gfp* (pHB6458) and a similar plasmid with TCCGGA replaced with GATATC (pHB6486). GFP was expressed in (pro)heterocysts of *Anabaena* 7120 [pHB6458] but not in differentiating cells of 7120*hetZ*del4-201 carrying the same plasmid; substitutions at TCCGGA abolished the expression of *gfp* in the wild-type strain (Fig. 2B). These results established that activation of *patS* in (pro)heterocysts largely depends on HetZ and the DIF1-motif promoter. Similarly, expression from the DIF1-motif promoter of *patX* is also dependent on the function of *hetZ* (S3 Fig).

### Upregulation of *hetR* in heterocysts depends on the DIF1 motif and *hetZ*

As shown with RT-qPCR, *hetR* was upregulated in the 7120*hetZ*del4-201 strain at 6 h after nitrogen stepdown (S4 Fig). However, the expression of *hetR* in *hetZ* mutants was probably not patterned [27].

*hetR* is an autoregulated gene [40], and a potential HetR-binding site has been identified upstream of the tsp −271 (for heterocyst-specific expression) [23, 26]. Upstream of the same tsp, there is also a potential DIF1-motif promoter [33]. To clarify the role of the HetR-binding site and the DIF1 motif in expression of *hetR*, we compared the expression of *gfp* from the promoter (−695 ~ −250 relative to the translational start site) of *hetR* and the same DNA fragment without the HetR-binding site or the DIF1 motif. Expression from the promoter of *hetR* was upregulated in (pro)heterocysts of *Anabaena* 7120, and the upregulated expression was abolished by substitutions at the DIF1 motif but not at the HetR-binding site (Fig. 3).

**Fig 3.**
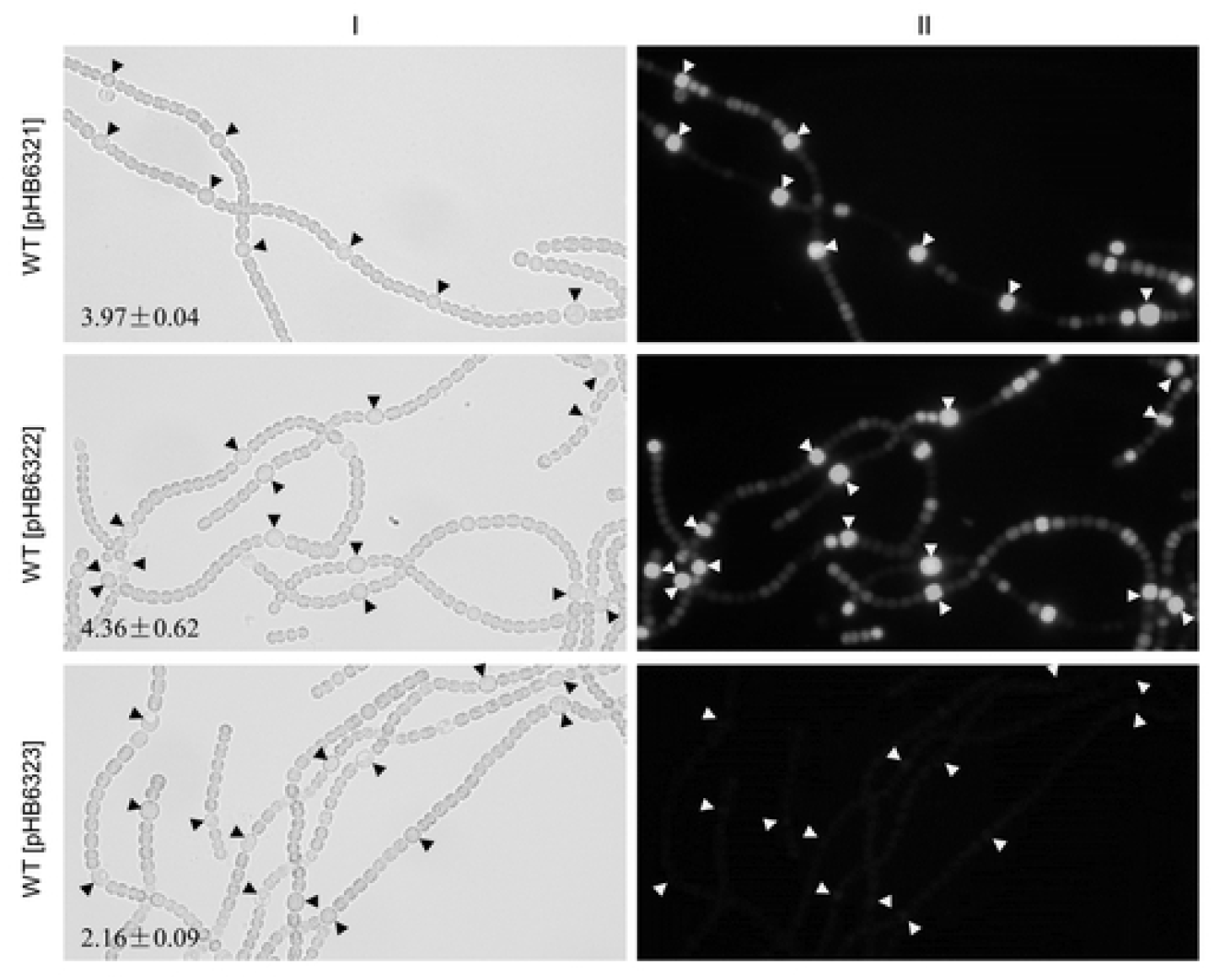
Light (I) and GFP fluorescence (II) photomicrographs showing the expression of *gfp* from the wild type or mutated promoter of *hetR* in *Anabaena* 7120. pHB6321: with the wild type promoter (−695 ~ −250) of *hetR;* pHB6322: with GGGN_5_CCC (potential HetR-binding site) in the promoter of *hetR* substituted with AAAN_5_TTT; pHB6323: with TCCGGA (DIF1 motif) in the promoter of *hetR* substituted with CAATTG. Solid arrowheads point to heterocysts; means ± SD are relative copy numbers of plasmids.

To confirm the role of the DIF1 motif in heterocyst-specific expression of *hetR*, we constructed a zeta-based plasmid with the minimal DIF1-motif promoter (a 40-bp fragment) upstream of *gfp* (pHB6821) and introduced the plasmid into *Anabaena* 7120 and the *hetZ* mutant. As shown in Fig. 4, GFP was expressed in (pro)heterocysts in *Anabaena* 7120 [pHB6821] but barely expressed in differentiating cells of the *hetZ* mutant. The copy numbers of zeta-based plasmids showed some changes in different strains but were still comparable in those in the same figure. Apparently, the upregulated expression of *hetR* in (pro)heterocysts is also mediated by HetZ via the DIF1 motif promoter.

**Fig 4.**
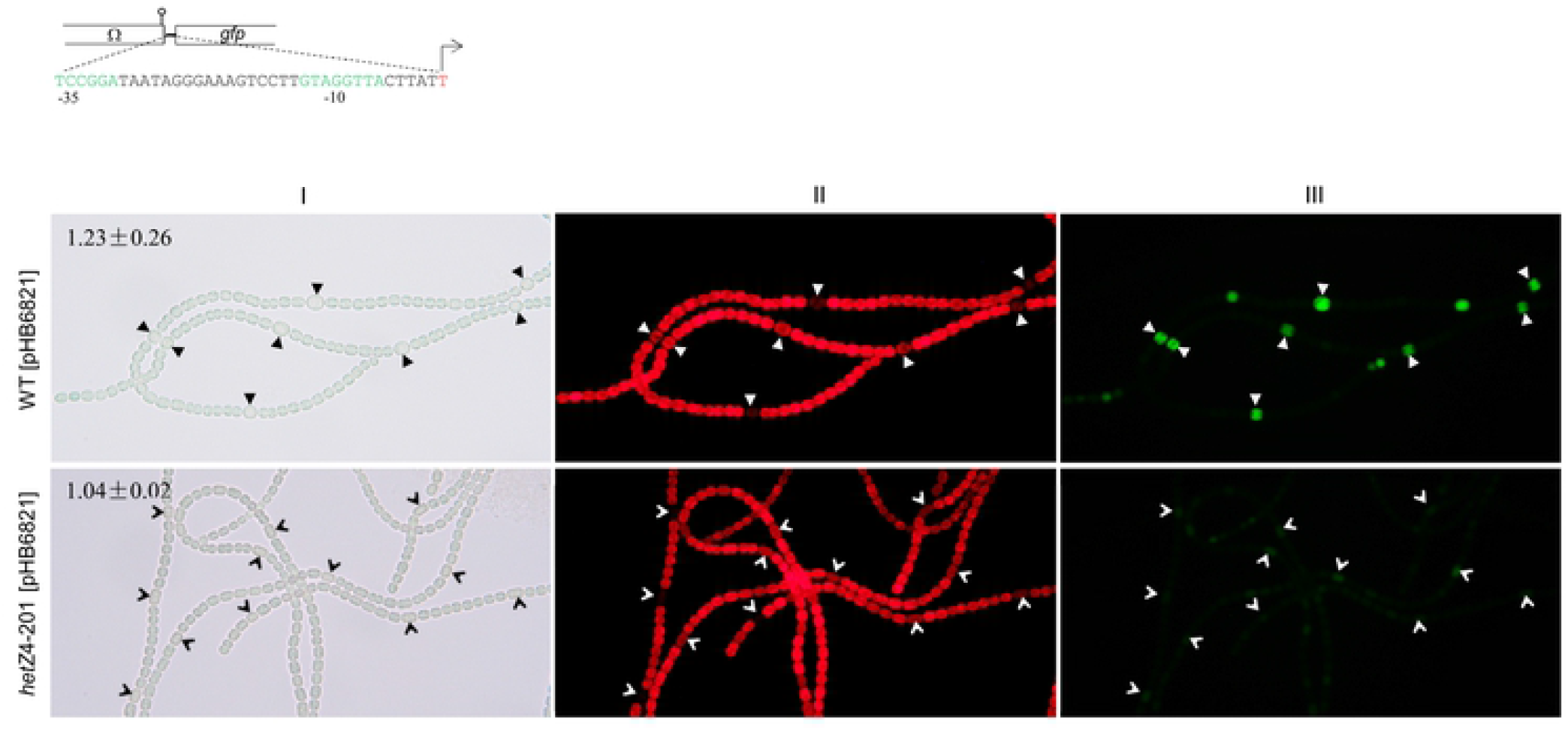
Expression of *gfp* from the minimal DIF1-motif promoter of *hetR* on a zeta-based plasmid in *Anabaena* 7120 and 7120*hetZ*del4-201. Top: the minimal sequence of DIF1-motif promoter cloned upstream of *gfp* in pHB6821. Photographs: light (I), autofluorescence (II) and GFP fluorescence (III) photomicrographs of *Anabaena* 7120 and the *hetZ* mutant with pHB6821 at 24 h after nitrogen stepdown. Solid and empty arrowheads point to heterocysts and differentiating cells; relative copy numbers of plasmids are indicated as means ± SD.

We further generated a mutant of *Anabaena* 7120, P*_hetR_*-DIF1^-^, with the DIF1 motif substituted with GATATC in the chromosomal DNA. Compared to the wild type, the P*_hetR_*-DIF1^-^ strain showed delayed heterocyst differentiation and lowered heterocyst frequency (Fig. 5). Using RACE-PCR, we confirmed that the tsp at nucleotide −272 (−271 in previous reports [41, 42]) upstream of *hetR* in the wild-type strain disappeared in P*_hetR_*-DIF1^-^. Clearly, the DIF1 motif is required for the heterocyst-specific expression of *hetR* and normal heterocyst differentiation.

**Fig 5.**
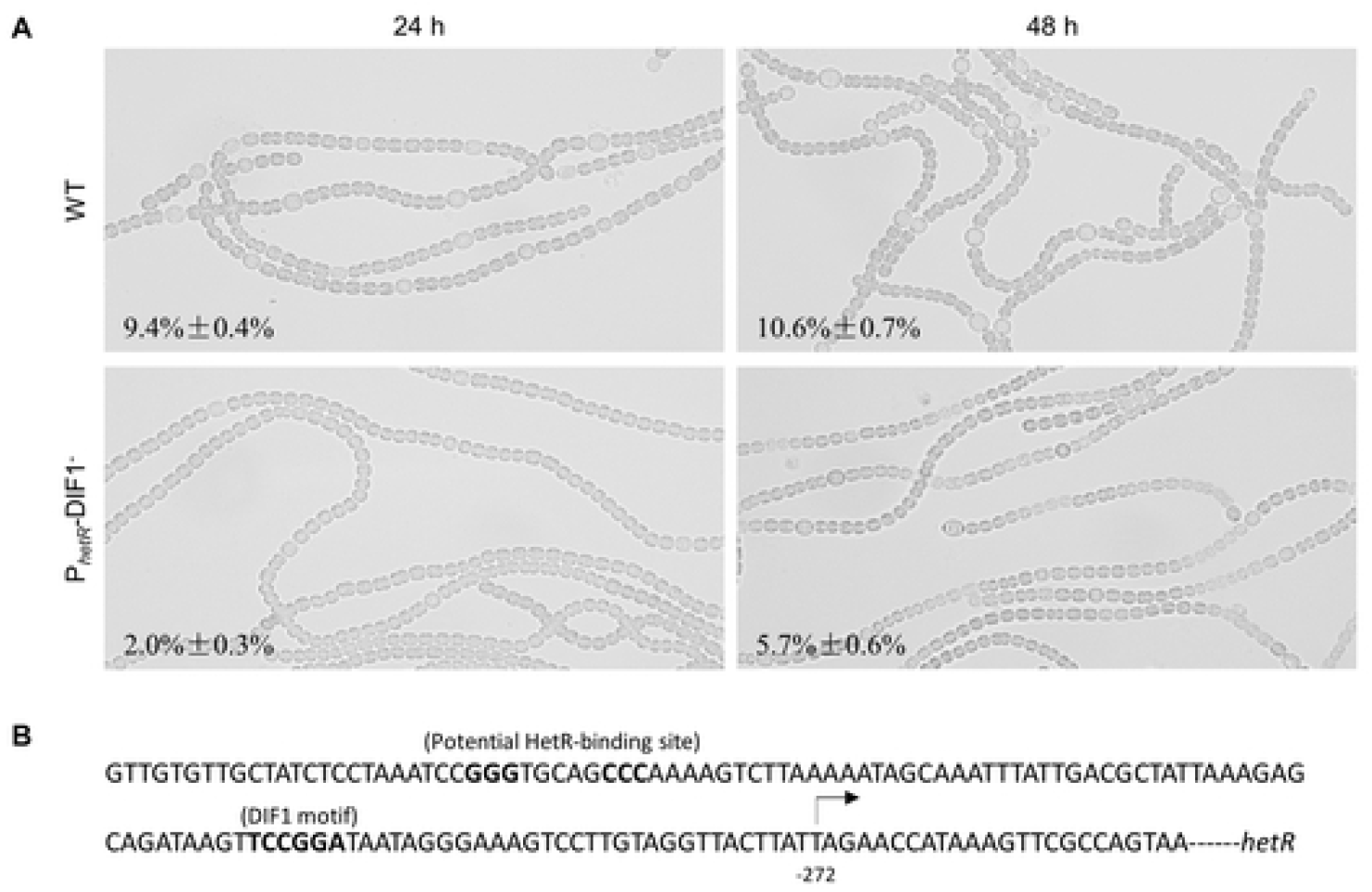
Differences between *Anabaena* 7120 and the P_*hetR*_-DIF1^-^ strain in heterocyst differentiation and expression of *hetR*. (A) Photomicrographs of *Anabaena* 7120 and the P_*hetR*_-DIF1^-^ strain at 24 h and 48 h after nitrogen stepdown. Frequencies of heterocysts/proheterocysts are indicated. (B) A stretch of sequence upstream of *hetR* showing the DIF1 motif, potential HetR-binding sequence and the tsp at −272.

### PatU3 interacts with HetZ and modulates the expression of *patS* and *hetR*

*hetZ* and *patU3* play opposite roles in heterocyst differentiation, whereas *patU5* (which lies between *hetZ* and *patU3*) is not involved in heterocyst differentiation [27]. Employing the yeast two-hybrid system, we found that PatU3 interacts with HetZ (Fig. 6A-i); by a pull-down experiment, we confirmed the interaction between the two proteins (Fig. 6B). As indicated in the two-hybrid assay, HetZ without the C-terminal portion no longer interacted with PatU3 (Fig. 6A-ii).

**Fig 6.**
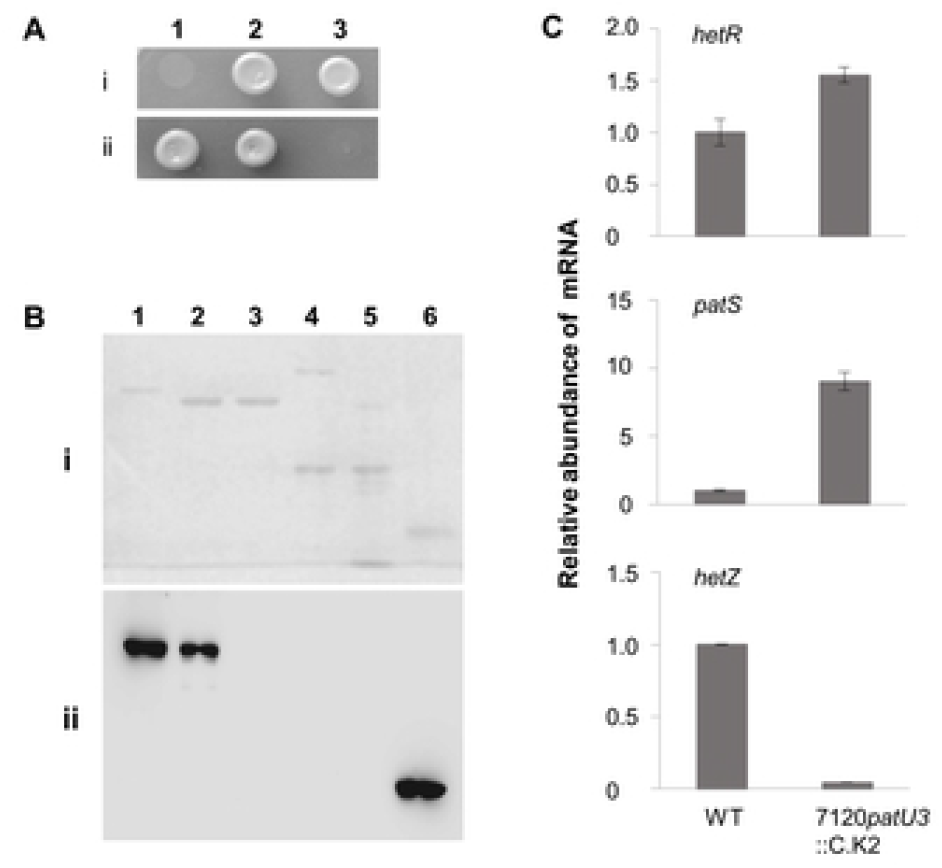
Interaction of PatU3 with HetZ. (A) Yeast two-hybrid assays of the interaction between PatU3 and HetZ. i) 1, pGBKT7-Lam + pGADT7-T, as the negative control; 2, pGBKT7-53 + pGADT7-T, as the positive control; 3, pGBKT7-PatU3 + pGADT7-HetZ. ii) 1, pGBKT7-PatU3 + pGADT7-HetZ[2-144]; 2, pGBKT7-PatU3 + pGADT7-HetZ[145-288]; 3, pGBKT7-PatU3 + pGADT7-HetZ[289-401]. Bracketed numbers (amino acid residue no.) indicate the portion deleted from HetZ (which has a full length of 401 aa). (B) Pull-down assays of the interaction. Proteins were separated by SDS-PAGE (I) and analyzed with Western blot detection using anti-HA monoclonal antibody (II). 1, EF-Ts(HA)-HetZ; 2, MBP-PatU3 + MBP·Bind resin + EF-Ts(HA)-HetZ; 3, MBP-PatU3 + MBP·Bind resin + EF-Ts(HA); 4, MBP + MBP·Bind resin + EF-Ts(HA)-HetZ; 5, MBP + MBP·Bind resin + EF-Ts(HA); 6, EF-Ts(HA). (C) RT-qPCR analysis of mRNA abundance of *hetR, patS* and *hetZ* in *Anabaena* 7120 and the *patU3*::C.K4 mutant at 6 h after nitrogen stepdown. Data are means ± SD of 3 technical replicates.

The interaction between PatU3 and HetZ may modulate HetZ-dependent gene expression. Based on RT-qPCR analysis, we compared the expression of *hetR* and *patS* in the wild type and the 7120*patU3*::C.K4 strain at 6 h after nitrogen stepdown (Fig. 6C). Relative to the wild type level, the mRNA level of *patS* was greatly increased in the *patU3* mutant, whereas that of *hetR* was slightly increased. Increased expression of *patS* probably inhibited the transcription of *hetZ* in the mutant (P*_hetZ_-gfp* in the mutant had shown a similar result, see ref. 27). However, the *patU3*::C.K4 mutation did not change the abundance of proteins HetR and HetZ in *Anabaena* filaments (S5 Fig).

## Discussion

HetR and PatS-derived peptides are key players for heterocyst differentiation and patterning in *Anabaena* 7120. How their encoding genes are regulated is an important question for understanding the molecular mechanism of the differentiation/patterning process. In this study, we show that the DIF1 motif plays an important role in regulation of these genes and that expression from DIF1 promoters depends on the function of *hetZ*.

HetR is often considered as the master regulator of heterocyst differentiation, and it directly regulates the expression of *hetP* [21] and *hetZ* [23] in developing heterocysts via HetR-recognition sequences and is required for the expression of *patA* in vegetative cells [26]. How HetR controls the expression of *patS* and its own gene has not been clarified. By examining gene expression in *hetR*-minus heterocysts, we were able to show that HetR is non-essential for the upregulated expression from promoters of *hetR* and *patS* during heterocyst differentiation. Therefore, HetR may control the expression of these genes through other regulatory factors.

In sequences upstream of *hetR*, *patS* and *patX*, there are predicted DIF1-motif promoters. Synthetic minimal promoters of these genes all showed upregulated expression during heterocyst differentiation. Substitutions at the DIF1 motif greatly reduced the transcription activity of P*_patS_*. A mutant of *Anabaena* 7120 with the DIF1 motif of *hetR* substituted in the genome showed no transcription from the tsp −272 (or −271), which would otherwise be specifically activated in developing heterocysts [41]. Upstream of *hetR*, there is also a potential HetR-recognition site, but that site was shown to be not required for the upregulated expression. These results provided experimental evidence for the role of DIF1-motif promoters in heterocyst-specific expression of *hetR, patS* and *patX*.

On a plasmid with *gfp* as the reporter gene, minimal DIF1-motif promoters for *hetR*, *patS* and *patX* showed greatly weakened transcription in the 7120 *hetZ*del4-201 strain compared to those in the wild type. Expression of *gfp* from the full-length promoter of *patS* in the chromosome produced a similar result. Therefore, *hetZ* is required for the expression of DIF1-motif promoters.

Videau et al. showed that deletion of *hetZ* blocked the induced expression of *hetR, hetP* and *patS* [30]. A similar effect of *hetZ* mutation on the expression of *patS* had been shown in our previous study [27]. These observations could be explained as dependence of the expression of *patS* on either HetR or HetZ or both. In this study, we showed that HetR is not essential for the heterocyst-specific expression of *patS* and that HetZ is more directly involved in regulation of *patS*. For *hetR*, we found that the DIF1 motif rather than the HetR-binding site is required for the heterocyst-specific expression.

For the results we presented, two points need to be addressed in particular. (1) HetR and the global nitrogen regulator NtcA are dependent on each other for upregulated expression during heterocyst differentiation [43], how to explain the upregulation of P*_hetR_* in a *hetR*-minus background? There is no evidence that NtcA and HetR directly regulates each other. In at least one substrain of *Anabaena* 7120, NrrA mediates the regulation of *hetR* by NtcA [44, 45]. Proteins that mediate the regulation of *ntcA* by HetR have not been identified. Formation of functional heterocysts in the *hetR* mutant with P*_ntcA_-hetZ-hetP* implies that genes regulated by NtcA are properly expressed in developing cells. Presumptively, the expression of *hetZ* and *hetP* from P*_ntcA_* allowed sufficient expression of NtcA in developing cells, and NtcA in turn enhances the expression of P*_ntcA_-hetZ-hetP* and indirectly upregulates P_*hetR*_. (2) How to explain the differentiating cells in the 7120del*hetZ*4-201 mutant? In this *hetZ* mutant generated with the substrain of *Anabaena* 7120 in our laboratory, we found that the mRNA level of *hetR* was increased after nitrogen stepdown as in the wild type (S4 Fig), even though the expression was probably not patterned. The expression of *hetR* can initiate cell differentiation (that ceases at the very early stage) in a less regular pattern (consistent with the low expression of *patS*).

In addition to HetZ, we also analyzed PatU3, an inhibitory protein factor for heterocyst differentiation. Protein interaction assays indicated that PatU3 interacts with HetZ. In a *patU3* mutant, the transcription of *patS* was greatly enhanced, that of *hetR* slightly enhanced, *hetZ* greatly inhibited, relative to the wild type levels; however, the abundance of proteins HetR and HetZ remained unchanged. PatU3 appeared to exert complicated effects on the expression of *hetR*, *hetZ* and *patS* at mRNA and protein levels. Presumptively, PatU3 can regulate the cellular concentration of free HetZ (relative to the PatU3-bound form) and affect the stability of HetZ, therefore modulate HetZ-dependent gene expression. However, PatU3 may also have additional functions that indirectly affect the expression of these genes.

As a gene directly regulated by HetR, *hetZ* is involved in initiation of heterocyst differentiation and regulation of *patS*/*patX* and *hetR*. Although there is no evidence that HetZ directly interacts with the DIF1 motif promoter, it is clear that HetZ is more direct than HetR in regulation of *patS/patX*. Therefore, HetR, HetZ and PatS/PatX form a molecular circuit of mutual regulation (but the role of *patX* in de novo heterocyst patterning awaits experimental investigation). This conclusion is important, because HetZ provides an additional site for modulation of the expression of *patS*/*patX*, which are the sources of diffusible inhibitor for de novo pattern formation at the early stage. PatU3 is a candidate for the modulator. It interacts with HetZ and somehow modulates the expression of *hetR, hetZ* and *patS*. This coordination scenario involving multiple activating/inhibiting factors may help to refine the current models [46, 47] for heterocyst differentiation and patterning.

## Acknowledgements

This work was supported by the National Natural Science Foundation of China (Grant numbers 31770044 and 31270132), the State Key Laboratory of Freshwater Ecology and Biotechnology at IHB, CAS (2019FBZ09) and the Knowledge Innovation Project of Hubei Province (2017CFA021).

## Supporting Information

**S1 Fig.** RT-qPCR analyses showing the role of *hetZ* in expression of *patS* during heterocyst differentiation.

**S2 Fig.** Expression of *gfp* fused to fragments upstream of *patS* on a pDU1-based plasmid in *Anabaena* 7120.

**S3 Fig.** Light (I), autofluorescence (II) and GFP fluorescence (III) photomicrographs showing the expression of *gfp* from the DIF1-motif promoter of P*_patX_* in *Anabaena* 7120 and 7120*hetZ*4-201.

**S4 Fig.** RT-qPCR analysis of the expression of *patS* and *hetR* in the wild type and the mutant 7120*hetZ* del4-201 at 0 and 6 h after nitrogen stepdown.

**S5 Fig.** Detection of HetR and HetZ in the wild type and the *patU3* mutant of *Anabaena* 7120 at 24 h after nitrogen stepdown.

**S1 Table.** *Anabaena* strains, plasmids and primers.

